# RNA splicing alterations induce a cellular stress response associated with poor prognosis in AML

**DOI:** 10.1101/2020.01.10.895714

**Authors:** Govardhan Anande, Nandan P. Deshpande, Sylvain Mareschal, Aarif M. N. Batcha, Henry R. Hampton, Tobias Herold, Soren Lehmann, Marc R. Wilkins, Jason W.H. Wong, Ashwin Unnikrishnan, John E. Pimanda

## Abstract

RNA splicing is a fundamental biological process that generates protein diversity from a finite set of genes. Recurrent somatic mutations of splicing factor genes are relatively uncommon in Acute Myeloid Leukemia (AML, < 20%). We examined whether RNA splicing differences exist in AML even in the absence of splicing factor mutations. Analyzing RNA-seq data from two independent cohorts of AML patients, we identified recurrent differential alternative splicing between patients with poor and good prognosis. These alternative splicing events occurred even in patients without any discernible splicing factor mutations. The alternative splicing events recurrently occurred in genes involved in specific molecular functions, primarily related to protein translation. Developing informatics tools to predict the functional impact of alternative splicing on the translated protein, we discovered that ~45% of the splicing events directly affected highly conserved protein domains. Several splicing factors were themselves misspliced in patients, and the splicing of their target transcripts were also altered. By studying differential gene expression in the same patients, we identified that alternative splicing of protein translation genes in ELN^Adv^ patients resulted in the induction of an integrated stress response and up-regulation of inflammation-related genes. Lastly, using machine learning techniques, we identified a set of four genes whose alternative splicing can refine the accuracy of existing risk prognosis schemes and validated it in a completely independent cohort. Our discoveries therefore identify aberrant alternative splicing as a molecular feature of adverse AML with clinical relevance.

**Key Points:**

- Widespread and recurrent alternative splicing differences exist between AML patients with good or poor prognosis
- Missplicing of RNA splicing factors leads to altered splicing of their target transcripts
- Aberrant splicing of protein translation genes triggers the induction of an integrated stress response and concomitant inflammatory response
- Alternative RNA splicing information can be used to improve the accuracy of existing prognostic algorithms in AML

## Introduction

Acute Myeloid Leukemia (AML) is a hematological malignancy associated with a poor prognosis and a <30% five-year survival rate ^1^. With an incidence rate of 4 per 100,000 adults per year ^2^ and a five-fold higher rate in people over the age of 65, AML represents ~40% of all new adult-onset leukaemias in developed societies ^3^. AML is characterized by the clonal proliferation of undifferentiated myeloid precursor cells in the bone marrow and impaired haematopoiesis ^4^. AML patients have recurrent somatic driver mutations ^5–7^ in addition to characteristic cytogenetic and chromosomal abnormalities. These alterations have prognostic significance and are used to classify AML ^5^. However, not all of these mutations are exclusive to AML, with many also being detected in Myelodysplastic Syndrome (MDS) ^8,9^ as well as in healthy individuals with age-related clonal haematopoiesis ^10,11^.

The standard of care treatment for AML is intensive induction chemotherapy. However, despite complete remission (CR) rates of >50%, long term disease-free survival remains poor at <10% and a median overall survival of less than 12 months in patients aged over 60 years ^12^. Additionally, because of significant co-morbidities, intensive chemotherapy may not suit older patients ^13,14^. Alternate therapies for these individuals may include lower-intensity treatments, DNA hypomethylating agents (HMA) ^15,16^ or targeted therapies. However, response rates and survival benefits still remain poor ^17^, highlighting an important need to develop new therapeutic options for the management of AML.

In order to develop more effective drugs for AML, it is necessary to better understand the molecular aberrations present in leukemic cells. Aberrations in RNA splicing, a fundamental and highly conserved process occurring in >95% of multi-exon human genes ^18^, are increasingly being described in many cancers ^19^. Splicing is a co-transcriptional event, orchestrated by *cis*-acting regulatory elements as well as *trans*-acting factors of the spliceosomal complex ^20^. Pan-cancer studies have begun to reveal that tumors have an average of ~20% more alternative splicing events than in healthy samples ^19,21^. Dysregulation of the expression of splicing factors ^22,23^ and upstream signaling pathways ^24^ as well as genomic mutations in *cis*-splice sites ^25,26^ have all been reported in cancers. Additionally, in hematological malignancies such as MDS and Chronic Myelomonocytic Leukemia, recurrent somatic mutations in members of the E- and A-splicing complexes, such as SF3B1, U2AF1, SRSF2 and ZRSR2 are detected in >50% of patients ^8,9,27^. The exact mechanisms through which these mutations contribute to the malignancy remain poorly understood. *U2AF1* and *SF3B1* mutations might alter the 3’ splice site in target transcripts ^28–30^, *SRSF2* hotspot mutations affect the preferred binding motif on transcripts ^31–33^ while *ZRSR2* gain-of-function mutations increase intron retention ^34^. The different mutations have been proposed to affect unique sets of genes ^29,33^ although convergence has been proposed at the level of pathways ^35^ or through the induction of R-loops ^36^.

Somatic mutations in the splicing machinery are less frequent in AML however. Analyses of large cohorts of patients have determined the overall frequency of splicing mutations to be <20% ^6,7^. However, widespread dysregulation of RNA splicing has been observed even in cancers with low frequencies of splicing factor mutations ^19,21,37^. We therefore examined whether RNA splicing alterations exist in AML even in the absence of somatic splicing factor mutations, and whether it correlates with disease outcome.

## Methods

### Patient cohorts

Data from two adult AML cohorts were used in the discovery phase of the study: The Cancer Genome Atlas (TCGA)-AML cohort ^38^ and the Clinseq-AML cohort ^39^. Data from the Beat-AML cohort ^7^ was used to validate significance of the splicing signature. Full details are provided as Supplemental Data.

### RNA-seq data analyses

RNA-seq data were analyzed for multiple types of alternative splicing using a custom in-house bioinformatics pipeline incorporating available tools, including Mixture of Isoforms (MISO) to determine Percent Spliced In (PSI) values in each sample and rMATS for differential splicing analyses. Differential gene expression analyses were performed using DESeq2. A custom in-house pipeline was developed to identify possible changes in well-annotated protein domains due to differentially spliced events. Full details are provided as Supplemental Data.

### Transcript motif analyses

Predictions of differential binding of RNA binding proteins were made using rMAPS ^40^. Maximum entropy modeling was done with MaxEntScan ^41^. Full details are provided as Supplemental Data.

### Prognostic model generation

The splicing signature was generated using LASSO Cox Regression with ten fold cross validation implemented in glmnet (R package v 2.0-16) ^42^. The splicing risk score for each patient was calculated from the regression coefficients. Performances of prognostic models were assessed by Harrel’s C index. Risk contributions and variable importance of all prognostic models were estimated as described previously ^43^. Full details are provided as Supplemental Data.

## Results

### Identification of differential alternative splicing related to outcome in AML patients

To determine whether RNA splicing alterations might be a factor in adverse outcomes in AML, we developed a bioinformatics pipeline to quantify differential alternative splicing in RNA-seq data. We first analyzed AML transcriptomes from The Cancer Genome Atlas (TCGA) ^6^. To detect whether splicing alterations can exist even in the absence of any somatic mutations in splicing factors, we focused our analyses on patients who did not have any splicing factor mutations using the available somatic mutation data ^6^. To validate the mutational data, we queried the RNA-seq data to detect the presence of known splicing factor hotspot mutations ^27^ in transcripts. We confirmed the mutation data in all patients identified to have splicing factor mutations and identified an additional sixteen patients with splicing factor hotspot mutations detectable in the RNA-seq data (fourteen with *SRSF2* and one each with *SF3B1* and *ZRSR2*). Our findings of TCGA-AML patients with unannotated splicing mutations is consistent with a recent independent report ^44^.

We stratified patients in the TCGA-AML using the widely used European LeukemiaNet (ELN) prognostic scheme ^1,4^ (Fig 1A) and restricted the analysis to patients who received intensive induction chemotherapy and for whom full clinical data was available (n=104, Supplementary Table 1). Performing differential splicing analyses between ELN^Fav^ and ELN^Adv^ (Fig 1B), studying the five different types of alternative splicing events (schematized in Fig 1C), we identified 1288 differentially spliced events (at FDR ≤0.05) in 910 genes (Fig 1D, Supplementary Table 2). A majority of the events involved the differential skipping (or retention) of exons preferentially in one set of patients (n=716, 55.5%, Fig 1D). Of these, 395 events involved preferential skipping of exons in ELN^Adv^ patients (Supplementary Fig 1A), with the remaining (n=321) associated with exon skipping in ELN^Fav^ patients. An example of differential exon usage was the skipping of exon 37 of *MYO9B* in TCGA-ELN^Fav^ patients (Fig 1E). Only reads spanning exon 36 – exon 38 were detected in ELN^Fav^ patients (representative examples, patients #2914 and #2955, Fig 1E). In ELN^Adv^ patients (representative patients #2855 and #2817, Fig 1E) however, there was an increase in the number of reads indicating the inclusion of exon 37 (128 and 41 reads joining exons 36 and 37, and 114 and 61 reads joining exons 37-38, in ELN^Adv^ patients #2855 and #2817 respectively, Fig 1E; compared to no reads in the ELN^Fav^ patients) and a concomitant decrease in reads spanning exon 36 and 38, skipping exon 37 entirely (29 and 24 reads respectively, compared to 63 and 46 in ELN^Fav^ patients #2914 and #2955, Fig 1E). A related phenomenon, of mutually exclusive exon usage, where adjacent exons are alternately used, contributed to 185 differential events (Fig 1D). The retention of introns was the next most prevalent class (n=201, 15.6%, Fig 1D) in TCGA-ELN^Adv^ patients, as seen in the representative example of the retention of intron 8 in *CDK10* (increased intron-specific reads and a decrease in exon-exon reads in ELN^Adv^ patients, Fig 1F). Additional examples of differential 3’ or 5’ splice site usages are shown in Supplementary Figs 1B and 1C respectively.

**Figure 1:**
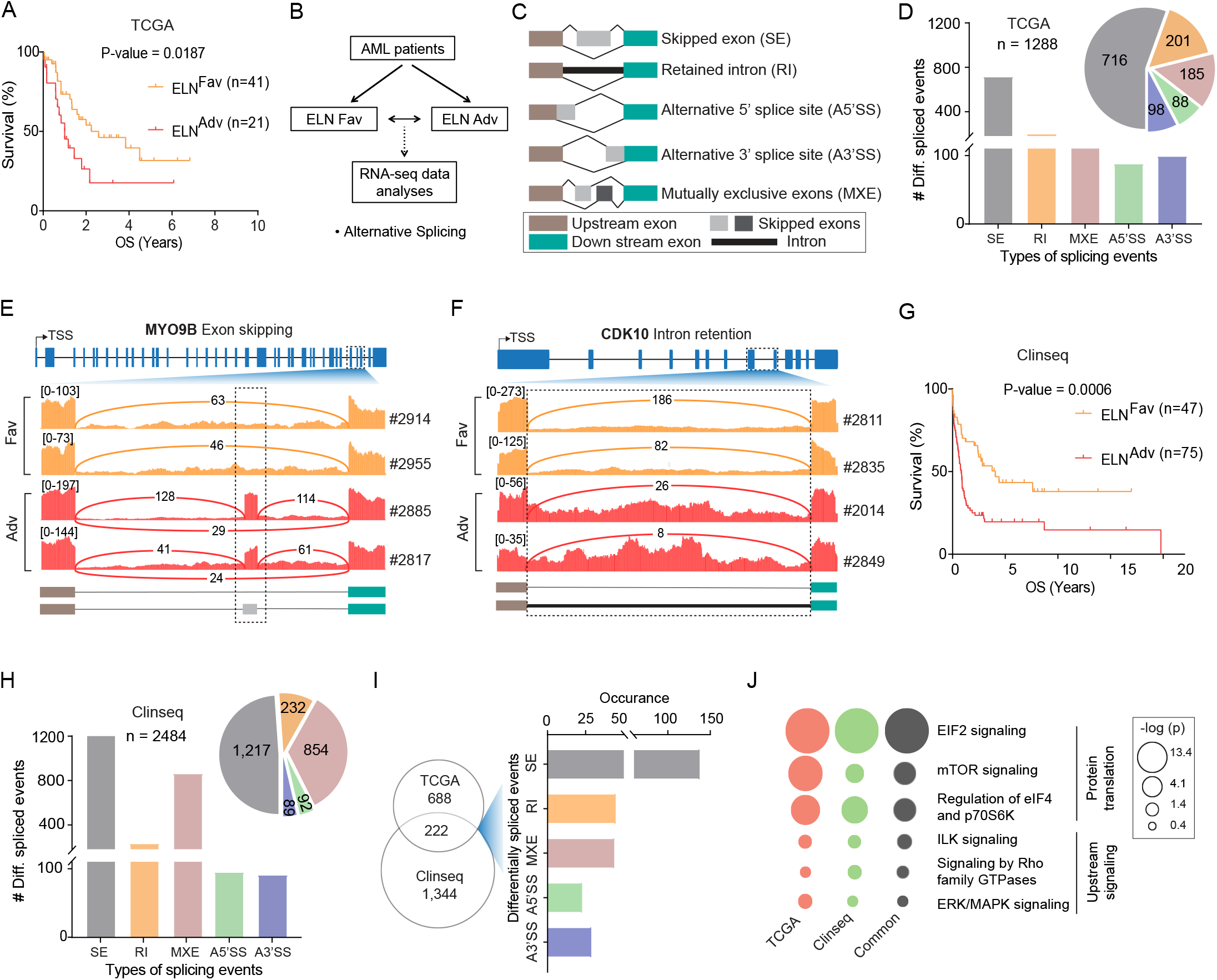
Identification of differential alternative splicing in AML patients. (A) Kaplan-Meier survival analysis of ELN stratification in the TCGA AML cohort showing significant survival differences between three ELN subgroups. P-value computed using Log-rank (Mantel-Cox) test. (B) Schematic outline to analyse RNA-seq data for differential alternative splicing. (C)Different alternative splicing events detected. Exons affected are represented in grey, while up- and down-stream exons are showed in brown and green respectively. Introns are represented as a black line, and depicted as a solid thick black line when retained. (D) Distribution of differentially spliced events identified comparing ELN^Fav^ and ELN^Adv^ in the TCGA AML cohort. SE, skipped exons. RI, retained introns. MXE, mutually exclusive exons. A5’SS, alternative 5’ splice sites. A3’SS, alternative 3’ splice sites. (E) Sashimi plot of a representative exon skipping event in the *MYO9B* gene in the TCGA data. Sequencing reads support the skipping of exon 37 (boxed) in two representative ELN^Fav^ patients (#2914, #2955, orange tracks), while indicating exon inclusion in ELN^Adv^ patients (#2855, #2817, red tracks). Lines connecting each exon represent splice junctions and numbers on each line represent number of supporting RNA-seq reads. (F) Sashimi plot of a representative intron retention event in the *CDK10* gene in the TCGA data. Retention of intron 8 (boxed) is observed in ELN^Adv^ patients (representative patients #2014, #2849, red tracks). Lines connecting each exon represent splice junctions and numbers on each line represent number of supporting RNA-seq reads. (G) Kaplan-Meier survival analysis of ELN stratification in the Clinseq AML cohort showing significant survival differences between three ELN subgroups. P-value computed using Log-rank (Mantel-Cox) test. (H) Distribution of differentially spliced events identified comparing ELN^Fav^ and ELN^Adv^ in the Clinseq AML cohort. SE, skipped exons. RI, retained introns. MXE, mutually exclusive exons. A5’SS, alternative 5’ splice sites. A3’SS, alternative 3’ splice sites. (I) Venn diagram depicting the overlap of differentially spliced genes in both cohorts. Bar plots represent the distribution of alternative splicing events in the shared set of genes. SE, skipped exons. RI, retained introns. MXE, mutually exclusive exons. A5’SS, alternative 5’ splice sites. A3’SS, alternative 3’ splice sites. (J) Bubble plots of Ingenuity Pathway Analysis of the differentially spliced genes, in TCGA (red), Clinseq (green) and shared genes (grey). The size of each bubble corresponds to significance of enrichment.

To validate these findings, we analyzed an independent cohort of AML patients from the Scandinavian Clinseq study ^39,45^. Selecting the patients similarly to the TCGA cohort, we performed differential splicing analyses between Clinseq-ELN^Fav^ (n=47) and ELN^Adv^ (n= 75) patients (Fig 1G). We detected a total of 2484 alternative splicing events (FDR ≤0.05), affecting 1566 genes (Fig 1H and Supplementary Table 3). As in the TCGA cohort, the majority of the events in the Clinseq data were skipped exons (n=1217, 48.9%, Fig 1H and Supplementary Fig 1A). Mutually exclusive exon usage was the next most prevalent (n=854, 34.4%) followed by intron retention (n=232, 9.25%, Fig 1H). Comparing both cohorts, we found differential splicing events occurring in the same direction, i.e. enriched either in ELN^Adv^ patients in both cohorts, or in ELN^Fav^ patients in both cohorts, in 222 genes (Fig 1I, Supplementary Table 4). Of these, 93 splicing events (in 78 genes) were identical in both cohorts, which we define as Class A events. A second class, Class B (244 events/173 genes in TCGA, 424 events/182 genes in Clinseq), affected the same gene and with the same directionality but represented different splicing events or occurred at different locations within the gene in the two cohorts. In 19 genes, we observed splicing occurring in opposite directions between the two cohorts.

To determine the molecular impact of this alternative splicing, we performed pathway analyses. Ingenuity Pathway Analysis of the Class A plus Class B genes revealed enrichment for a number of pathways, including those with functions related protein translation or intracellular signaling (Fig 1J and Supplementary Table 5). Orthogonal gene ontology-based analyses also supported these findings, with enrichment for pathways related to protein translation and RNA processing (Supplementary Fig 1D). Our data reveals recurrent and shared alternative splicing differences between AML patients with good or poor prognosis in two independent cohorts, converging on specific molecular pathways.

### Prediction of the functional consequences of alternative splicing

Analogous to genetic mutations, we expected that while some splicing events would have potentially deleterious effects on subsequent protein translation, others might be silent. To identify deleterious splicing events, we developed a custom bioinformatics pipeline (described in Methods). Briefly, the chromosomal coordinates of each splicing event were used to generate nucleotide sequences for the spliced and unspliced transcripts. These were then *in silico* translated and the generated primary sequences were scanned to predict the protein effect (Fig 2A).

**Figure 2:**
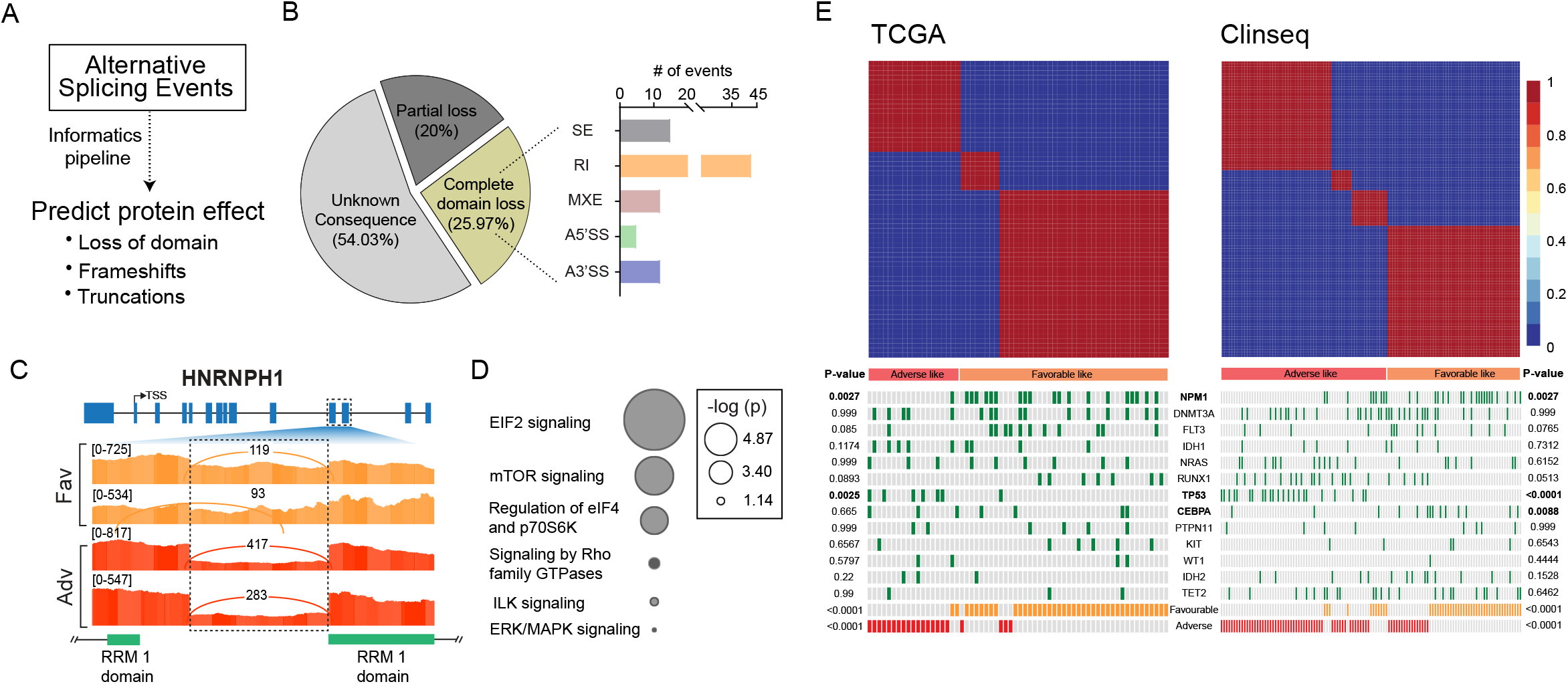
Analysis of the predicted impact of alternative splicing on protein function. (A) Schematic of the analytical pipeline to identify potentially deleterious alternative splicing events. (B) Pie chart distribution of protein domain prediction results. Bar plots on the right indicating the distribution of alternative splicing events predicted to lead to a complete loss of protein domains. (C) Sashimi plot of a representative protein domain disruption event caused by intron retention in *HNRNPH1* gene. Intron 11 is retained in ELN^Fav^ AML patients (two representative tracks shown, orange), disrupting the RRM1 domain. (D) Bubble plot of Ingenuity Pathway Analysis of genes with predicted complete loss of domains. The size of each bubble corresponds to significance of enrichment. (E) Non-negative matrix factorization clustering of AML patients based on similarity of splicing (Percent Spliced In (PSI) values) of the 222 shared splicing events, in the TCGA (left) and Clinseq (right) cohorts. Patients were classified as “adverse-like” (red), or “favorable-like” (orange) based on clustering. Oncoprints below denote somatic mutations identified in the patients. P-values (Fisher’s exact test) are shown for TCGA (left) and Clinseq (right), with events with p<0.05 in bold.

Using this methodological framework, we predict that 26% of the Class A plus Class B events (n=87, Fig 2B) cause a complete loss of well-annotated protein domains. The majority of these events involved intron retention events that alter the reading frame (Fig 2B). An example of this is the retention of intron 8 of the splicing regulator *HNRPH1* disrupting the <u>R</u>NA <u>R</u>ecognition <u>M</u>otif (RRM) protein domain (Fig 2C) and an altered transcript predicted to trigger nonsense-mediated decay. An additional 20% of events (n=67) lead to a partial loss of protein domains (Fig 2B). The functional consequences of a partial loss of a domain are harder to predict *a priori* and likely to be protein-specific. Furthermore, of the remaining 181 events (~54%) of unknown consequence (Fig 2B), we cannot rule out that some may also affect protein function, through altering protein secondary structure or unannotated domains. Focusing on the events leading to a complete domain loss, pathway analysis revealed that proteins affected by aberrant splicing are still enriched for specific molecular functions, including protein translation (Fig 2D), which we previously observed (Fig 1J). Our results suggest that alternative splicing changes leading to predicted protein dysfunction in genes involved in protein translation recurrently occur in AML patients.

### Analysis of the upstream drivers of alternative splicing differences

We next sought to understand the underlying reasons for the splicing differences in these risk groups. We investigated whether any of the somatic driver mutations (in the absence of splicing factor mutations) are correlated with the alternative splicing differences. We clustered patients based on the similarity of their splicing of Class A and B events and assessed the enrichment for somatic mutations within the clusters (Fig 2E). Apart from *NPM1*, *TP53* and *CEBPA* mutations, which are intrinsic to the ELN classification algorithm, no other somatic driver mutation showed any statistically significant correlation with the splicing groups (Fig 2E).

Amongst the alternatively spliced genes, we observed there were RNA binding proteins including factors with known roles in RNA splicing (Fig 3A). There was a trend for a greater number of predicted domain-altering events in these factors in ELN^Adv^ AML patients compared to ELN^Fav^ patients (Fig 3B). In addition, these factors form a tightly inter-connected network with multiple known protein-protein interactions (Fig 3C), suggesting that missplicing of these factors could trigger a cascade of splicing alterations in AML patients. To find evidence for this, we performed motif-scanning analyses ^40^ of the differentially spliced transcripts to determine if they might be targets for the misspliced splicing factors. RNA maps produced by rMAPS ^40^ indicated that a significant number of exons differentially retained in ELN^Adv^ AML have a target motif for HNRNPA1 (Fig 3D). Our informatics pipeline prediction was that the detected alternative splicing in *HNRNPA1*, a multi-functional splicing regulator that is known to act as a splicing repressor ^46,47^, would produce a non-functional protein in ELN^Adv^ patients. Consistent with this, exon inclusion in ELN^Adv^ patients was higher within transcripts where it would normally bind and repress splicing (Fig 3D). Conversely, HNRNPC is a splicing factor whose function is to repress exon inclusion ^48^ and predicted to be non-functional in ELN^Fav^ AML. Consistent with our predictions, we observed enrichment for HNRNPC motifs flanking exons that were differentially retained in ELN^Fav^ AML patients (Fig 3E).

**Figure 3:**
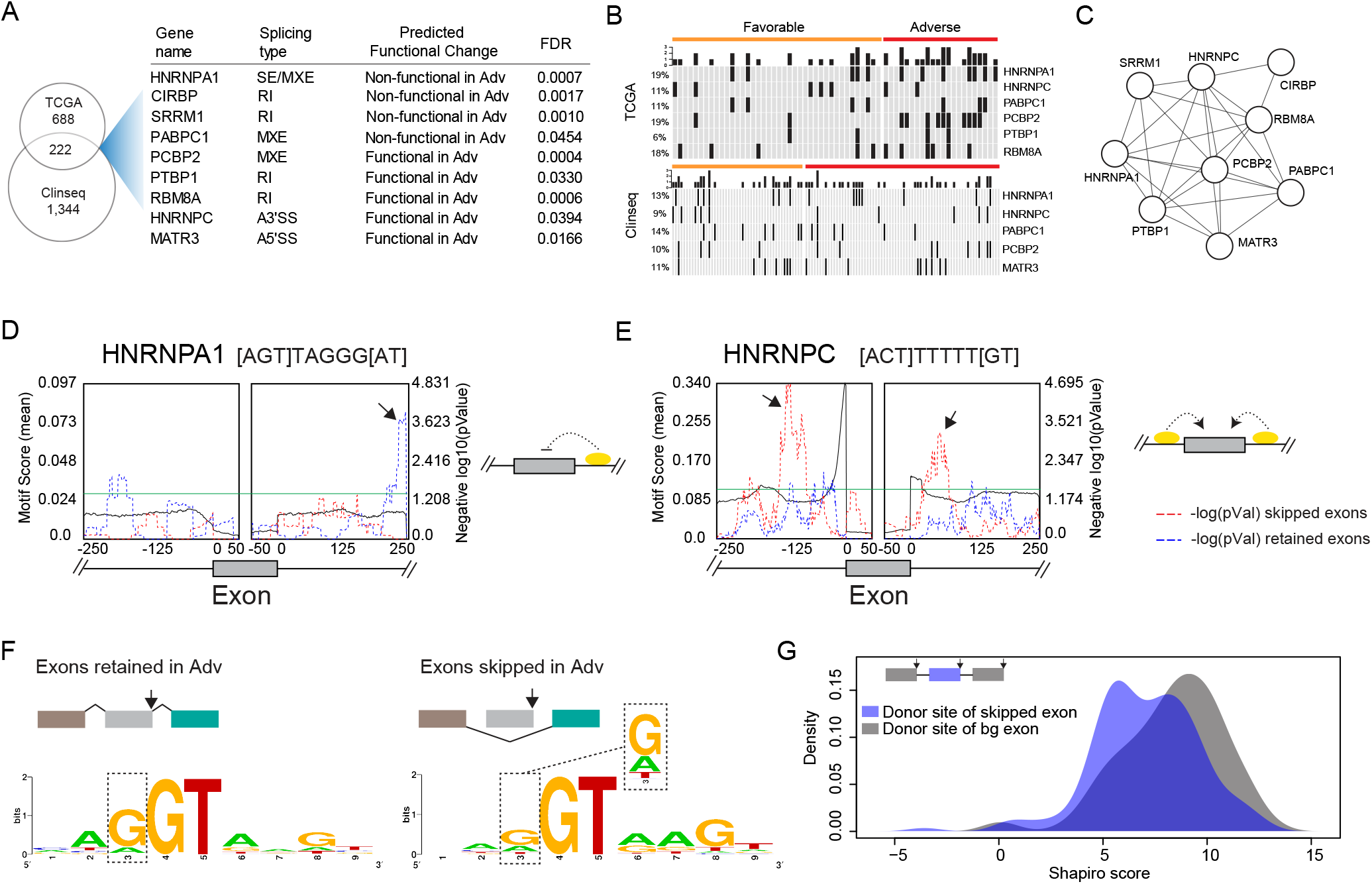
Analysis of the upstream drivers of differential alternative splicing in AML. (A) List of splicing factors that are commonly differentially spliced across both cohorts. The type of splicing, predicted effect on the protein and FDR are shown. (B) Plots indicating alternative splicing events, within splicing related genes (rows), predicted to have functional consequences on the translated protein in individual patients (columns) from the TCGA and Clinseq cohorts. ELN^Fav^ patients are on the left (yellow bar) and ELN^Adv^ patients on the right (red bar). (C) Interaction network indicating validated protein-protein interactions (edges) between the differentially spliced, with predicted functional impairment, splicing genes (nodes). (D) Motif scanning analysis for HNRNPA1 binding sites across a meta-exon generated from the differentially spliced events, with an arrow indicating a peak of significant over-enrichment. Motif enrichment scores (left axis) and P values (right axis) are shown. The dashed lines indicate scores of skipped (red) and retained (blue) exons, while the black solid line indicates that of a background score from all non-differentially spliced exons. The green horizontal line is set at p=0.05. (E) Motif scanning analysis for HNRNPC across a meta-exon generated from all differentially spliced events. Representation similar to (D). Arrows indicate peaks of significant over-enrichment. (F) LOGO analyses of splice donor sites of exons differentially retained (left) or skipped (right) in ELN^Adv^ patients. Analysis is within a 9-base window across the intron-exon junction (3 bases in exon and 6 bases in intron). (G) Smoothened density estimates of the position weight matrices (Shapiro score) of the splice donor sites of all differential exon-skipping events. Skipped exons (blue) and background exons (grey) are displayed, illustrating weaker splice sites in the skipped exons.

Extending motif analyses further, we found increased usage of non-canonical bases at the donor (Fig 3F) and acceptor (Supplementary Fig 2A) splice sites adjacent to exons differentially skipped in ELN^Adv^ patients. Furthermore, these skipped exons also have weaker donor sites ^49,50^ (Fig 3G). To determine whether dysregulation of other RNA binding factors (in addition to the ones we have already predicted above) might be contributing to differential splicing, we analyzed a catalogue of 114 well-characterized RNA binding motifs ^40^. We found enrichment of motifs for PABPC1, a RNA binding protein recently proposed to have roles in RNA splicing ^51^, and for RBM46 adjacent to exons that were retained in ELN^Adv^ patients (Supplementary Figs 2B-C). Similarly, introns that were preferentially retained in ELN^Adv^ were enriched for SRSF3 binding at the 3’ end (Supplementary Fig 2D). These RNA binding factors are also known to have protein-protein interactions with other splicing proteins predicted by our analyses to be affected by differential splicing (Supplementary Fig 2E). Our results suggest that missplicing of splicing factors, together with specific biophysical properties of *cis*-factors, contribute to the alternative splicing differences we have observed in AML patients.

### Induction of an integrated stress response in ELN^Adv^ AML patients

Our analyses had indicated that genes related to protein translation were differentially spliced (Fig 1J), with predicted functional impairment (Fig 2D) in ELN^Adv^ AML patients. A cellular consequence of impaired protein translation would be the induction of the <u>I</u>ntegrated <u>S</u>tress <u>R</u>esponse (ISR) within cells ^52^. To find evidence in support of this, we performed differential gene expression analyses between ELN^Fav^ and ELN^Adv^ patients (Fig 4A). 2219 genes were differentially expressed in the TCGA cohort at FDR <0.05 (Fig 4B, Supplementary Table 6), and 1710 genes in Clinseq (Fig 4B, Supplementary Table 6). GSEA analyses of the differentially expressed genes clearly indicate a strong up-regulation of ISR genes ^53^ in ELN^Adv^ AML patients in both cohorts (Fig 4C). Additionally, individual patient analyses revealed a proportional trend between the strength of the induction of ISR gene expression and the extent of missplicing of protein translation genes within the same patient (Fig 4D). ELN^Adv^ patients, who had higher levels of expression of ISR target genes, tended to have higher levels of missplicing of protein translation genes (Fig 4D).

**Figure 4:**
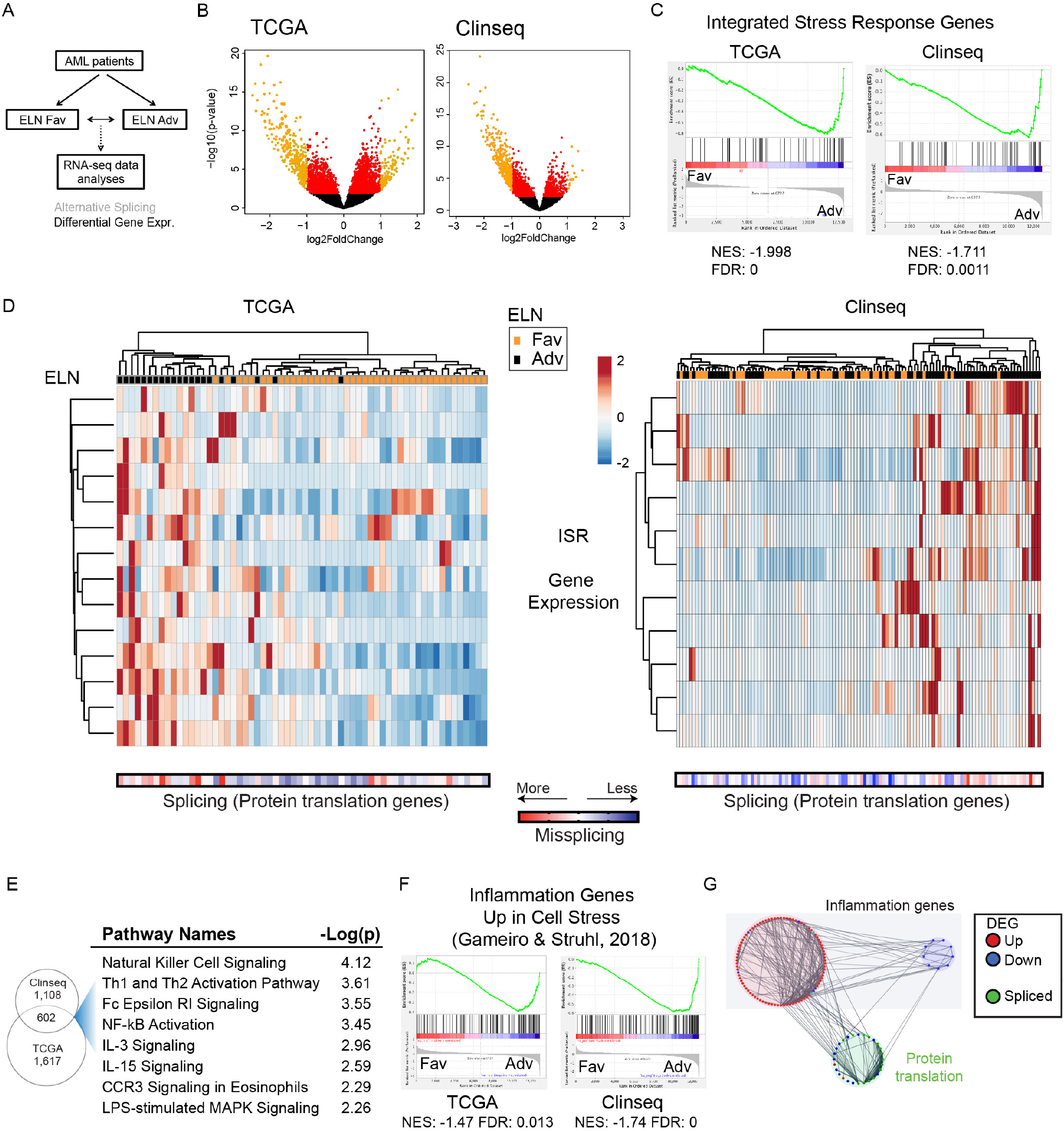
Analysis of the impact of alternative splicing on the transcriptome. (A) Schematic outline of the informatics methodology used to identify differentially expressed genes. (B) Volcano plots of differentially expressed genes in TCGA (left) and Clinseq (right). Genes highlighted in red have FDR < 0.05 and in orange with FDR < 0.05 and log_2_ (fold change) > |1|. (C) Gene set enrichment analyses (GSEA) showing up-regulation of integrated stress response genes ^53^ in ELN^Adv^ patients in TCGA (left) and Clinseq (right). (D) Hierarchical clustering of individual patients (columns) based on the expression of integrated stress response genes (rows) with core enrichment from GSEA for the TCGA and Clinseq cohorts respectively. Rows were scaled based on expression. A scaled z-score of the PSI values of protein translation genes was calculated in each patient and is represented below. (E) Ingenuity Pathway Analysis results of the differentially expressed genes. Venn diagram indicates differentially expressed genes shared by both cohorts. Inflammation-related pathways with associated enrichment values are shown. (F) GSEA analyses of a published ^54^ set of inflammation genes up-regulated as a result of decreased protein synthesis. Results show up-regulation in ELN^Adv^ patients in both the TCGA (left) and Clinseq (right) cohorts. (G) Integrated network analysis of differentially spliced translation genes (green) and differentially expressed inflammation genes (up-regulated in red, and down-regulated in blue). Experimentally validated protein-protein interactions are depicted as lines, connecting the proteins (nodes). DEG, differentially expressed genes.

It has recently been shown that metabolic stresses including amino acid deprivation that decrease protein synthesis trigger a pro-inflammatory response ^54^. Pathway analyses of the 602 genes that were commonly differentially expressed in the same direction in both cohorts (i.e. either up-regulated in ELN^Adv^ both cohorts, or down-regulated in both cohorts, Supplemental Table 5) revealed an enrichment for a number of inflammation-related pathways (Fig 4E). We find an up-regulation of these stress-induced inflammatory genes in ELN^Adv^ AML patients in whom protein translation is impacted due to splicing (Fig 4F). Network analyses further confirm strong interconnections between the misspliced translation-related genes and the differentially expressed pro-inflammation genes in ELN^Adv^ AML (Fig 4G). Our data supports a scenario where missplicing of protein translation genes trigger a pro-inflammatory stress response in ELN^Adv^ patients.

### Determining the prognostic relevance of alternative splicing events

Given these findings, we examined whether alternative splicing could serve as a prognostic marker for adverse outcomes in AML. While gene expression and epigenetic studies have been previously linked to AML outcome ^55–58^, these analyses would have missed the impact of alternative splicing. Utilizing machine-learning techniques (schematized in Fig 5A; see Supplementary methods for more details), we identified four genes (*MYO9B*, *GAS5*, *GIGYF2*, *RPS9*, Fig 5B) whose differential alternative splicing could stratify AML patients with good and poor prognosis. The differential splicing of these four genes (“*splicing signature*”) performs comparably to the ELN in both cohorts (Fig 5C), with similar Harrell’s C-statistic ^43,59^ (Supplementary Fig 3A) where a C-statistic of 50% is equivalent to a random assignment and 100% represents a correct ranking of the survival times of all patients.

**Figure 5:**
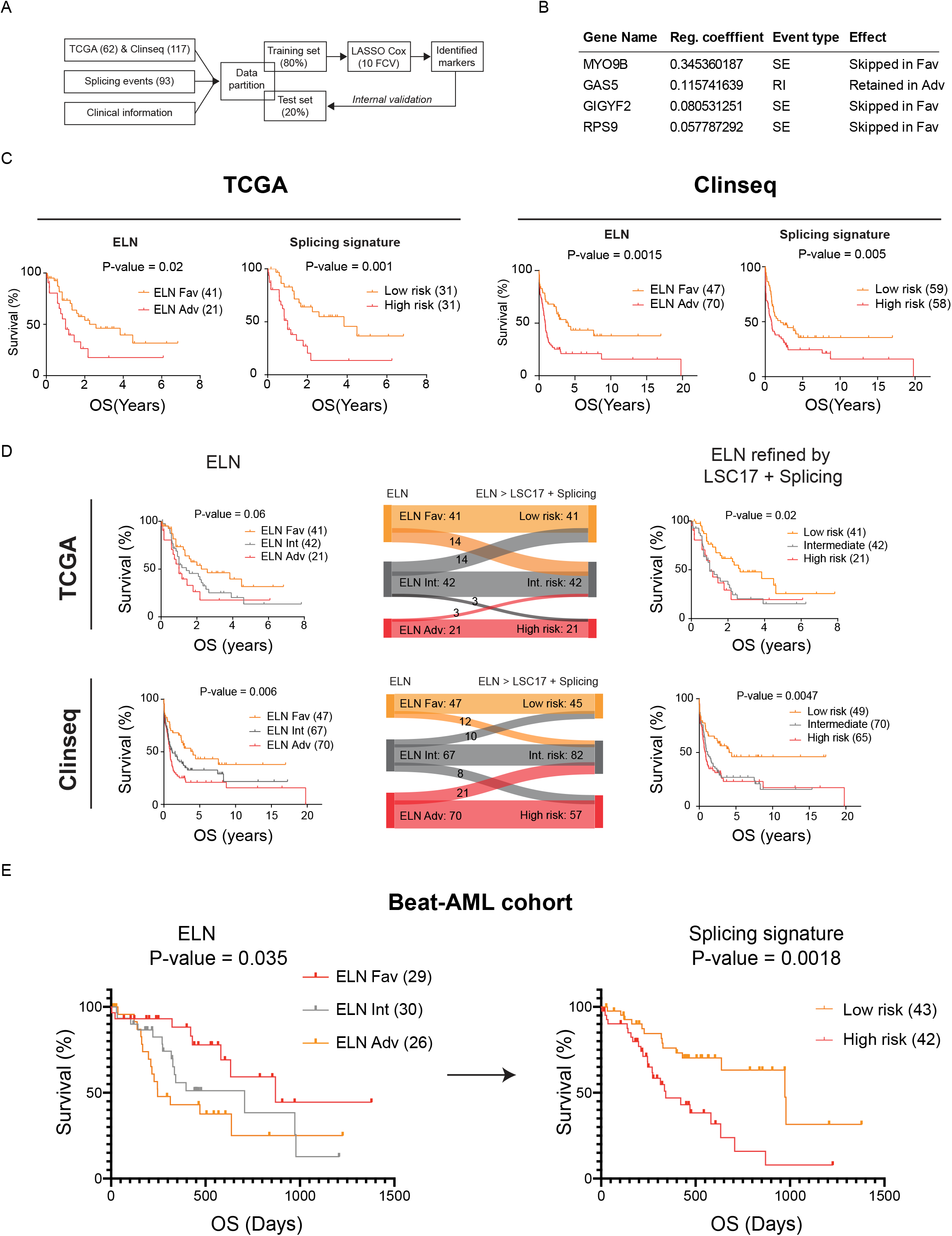
Evaluating the prognostic significance of alternatively splicing in AML. (A) Machine learning approach used to identify splicing markers. Input patients were randomly classified into two sets, training (80%) and test set (20%). LASSO cox regression with 10-fold cross validation was applied to the training set to identify markers. Identified markers were internally validated on the test set. (B) The four splicing markers identified to have prognostic significance in both AML cohorts. Regression coefficient and event type are displayed. (C) Kaplan-Meier analysis of TCGA-AML (top row) or Clinseq-AML patients (bottom row) stratified either by the ELN or the splicing signature. P-values were computed using Log-rank (Mantel-Cox) test. (D) TCGA-AML (top row) or Clinseq-AML patients (bottom row) classified initially by ELN (left panel) and re-classified by adding the LSC17 and the Splicing Signature (right panel). Sankey flow diagrams (middle panel) illustrate the redistribution of patients, with the widths of the lines proportional to numbers of patients redistributed (number also denoted). P-values were computed using Log-rank (Mantel-Cox) test. (E) Independent validation of the splicing signature in the Beat-AML ^7^ cohort. Patients were stratified by the splicing signature-based risk score. P-values were computed using Log-rank (Mantel-Cox) test.

More accurate stratification and improved prognosis would especially benefit AML patients classified as intermediate-risk, a group of patients with response and survival rates intermediate to ELN^Fav^ and ELN^Adv^. Accurately identifying ELN^Int^ patients with the most severe risk prognosis would aid in treatment decisions made in the clinic. Equally, ELN^Int^ patients with a predicted favorable prognosis could be treated appropriately. As the mutations and cytogenetics-based ELN, gene expression based LSC17 signature ^58^ and our splicing signature represent complementary biological measurements all with the potential to contribute to disease severity, we investigated their combined potential to more accurately classify AML patients. Addition of the splicing signature to the ELN or LSC17 alone improved the accuracy in both the TCGA (Supplementary Figs 3B-C) and Clinseq cohorts (Supplementary Figs 3D-E), with higher C-statistics for the combined signatures (Supplementary Figs 3F-G). Applied together, the combination of the three signatures improved the accuracy of classification of AML patients, converting the three-group risk classification to essentially two groups with significantly different overall survival in both cohorts (Fig 5D).

To independently validate the prognostic significance of the splicing signature, we also analyzed data from the BEAT-AML cohort ^7^. Selecting patients as for the TCGA and Clinseq cohorts, we performed RNA splicing analyses on the transcriptomic data and calculated the splicing risk score (see Supplementary methods for details). Applying the splicing signature significantly improved ELN 2017 based prognostic classification (p = 0.0018 vs p= 0.035, Fig 5E), converting the three-group risk classification into one with two groups with significantly different overall survival.

## Discussion

Recurrent somatic mutations in RNA splicing factors have been reported in several hematological malignancies ^60^. Analyzing AML transcriptomes, we have discovered recurrent alternative RNA splicing differences between ELN^Fav^ and ELN^Adv^ patients even in the absence of splicing factor mutations. Many of these alternatively spliced events are predicted to alter protein function, including members of the spliceosomal complex and protein translation genes. Integration with gene expression revealed that ELN^Adv^ patients had an induction of the ISR and a pro-inflammatory transcriptional program that was proportional to the degree of missplicing of protein translation genes. Furthermore, using machine learning, we identified four alternatively spliced genes that could be used to refine current mutation and transcriptome based prognostic classification of AML patients.

The origin of the missplicing that we have detected in AML patients remains unknown. It is possible that aberrant transcriptional programs initiated by oncogenic driver mutations might dysregulate splicing networks through the mis-expression of splicing co-factors. The splicing factors are also subject to a number of regulatory post-translational modifications. Phosphorylation of splicing factors by kinases of the SRPK and CLK families control their enzymatic activity and subcellular localization ^61^ and AML cells are sensitive to pharmacological inhibition of these kinases ^62^. Many RNA binding proteins are also methylated by the PRMT family of protein arginine methyltransferases and PRMT inhibition kills leukemic cells ^63^. Furthermore, splicing alterations due to epigenetic or chromatin changes due to somatic mutations ^44,64^ or possibly as a consequence of aging ^65^ have also been recently reported. It is possible that some or all of these mechanisms could contribute to the splicing alterations we have detected in AML. A cascade of missplicing would then be predicted to arise because of the highly-interconnected regulatory networks involving a number of splicing factors and RNA binding proteins ^32,46^.

Decreased protein translation induces the ISR, a conserved pathway which serves promotes cell survival by modulating cellular homeostasis during cellular stress ^52^. Protein translational stress leads to the efficient translation of the ISR effector *ATF4* and upregulation of its target genes ^52^. Increased ISR and ATF4 activity have been recently shown to be marker of leukaemic stem cells in AML patients ^53^. Our data indicates aberrant alternative splicing of protein translation genes and an induction of the ISR in AML patients with poor outcomes. Recently, a second cellular stress response –induction of a pro-inflammatory transcriptional program – has been identified as a result of decreased protein synthesis ^54^. Our data are consistent with this, where upregulation of inflammatory genes is seen in ELN^Adv^ patients.

Induction of inflammatory genes and the NFkB pathway have also been reported as a consequence of *SF3B1* and *SRSF2* mutations in MDS ^66^. It is possible that the functional consequences of aberrant RNA splicing, through somatic mutations or otherwise, might converge on common downstream consequences. The up-regulation of inflammation could induce a leukemic microenvironment that supports the growth of AML clones. AML cells have been recently reported to be dependent on signaling from the pro-inflammatory cytokine interleukin-1^67^. Furthermore, IL-1 signaling suppressed the growth of healthy leukemic cells, thereby promoting leukemogenesis and influencing clonal selection of neoplastic cells ^67^. While pharmacological inhibition of splicing factors has been proposed as a targetable vulnerability of leukemic cells ^65,68–71^, the narrow therapeutic window for these drugs due to toxicity poses a potential challenge to using them clinically. Our data suggests that targeting integrated stress response or inflammation-promoting pathways that might be stimulated in leukemic cells as a consequence of missplicing could be an alternative approach.

While cytogenetic and mutational information have become the clinical standard for prognosis in AML, there is still significant heterogeneity that remains unresolved. Assessing additional molecular parameters, including gene expression ^56,58^ and DNA methylation ^55^ have improved stratification of patients. However, these analyses would have missed capturing an important molecular feature of AML, aberrant alternative splicing. By complementing existing schema with splicing information, we were able to improve the accuracy of risk stratification, including for ELN^Int^ patients, which should aid in treatment decisions in the clinic.

## Acknowledgments

The authors would like to thank Dr. Ling Zhong (Mark Wainwright Analytical Centre, University of New South Wales) for technical assistance rendered. The authors also thank Dr. Annatina Schnegg (University of New South Wales) for critical review and discussion of the manuscript. This work was facilitated by infrastructure provided by the NSW Government co-investment in the National Collaborative Research Infrastructure Scheme (NCRIS, Australia).

The authors acknowledge the following funding support: GA was supported by a postgraduate scholarship from the University of New South Wales, with additional funding from the Translational Cancer Research Network. AU acknowledges funding support from the National Health and Medical Research Council of Australia (APP1163815), Leukemia & Lymphoma Society (USA) and Anthony Rothe Memorial Trust. JEP acknowledges funding from National Health and Medical Research Council of Australia (APP1024364, 1043934, 1102589), Cancer Institute of New South Wales/Translational Cancer Research Network and Anthony Rothe Memorial Trust. TH is supported by a grant of the Wilhelm-Sander-Stiftung (20130862) and the Physician Scientists Grant (G-509200-004) from the Helmholtz Zentrum München to TH and the German Cancer Consortium (Deutsches Konsortium für Translationale Krebsforschung, Heidelberg, Germany). TH is supported by a grant from Deutsche Forschungsgemeinschaft (DFG SFB 1243). AMNB is partially funded by the BMBF grant 01ZZ1804B (DIFUTURE).

## Authorship Contributions

The project was conceived by AU and the study design and experiments were planned by AU, NPD, JWWH and JEP. Most of the experiments and data analyses were performed by GA, guided and supervised by AU, NPD, JWWH and JEP. NPD, SM, AMNB and HH performed additional experiments and data analyses, with further supervision provided by TH, SL and MRW. Access to clinical information and transcriptomic data were generously provided by SL and TH. The manuscript was written by GA, NPD, AU and JEP, and reviewed and agreed by all coauthors.

