## Supplementary Methods, Suppl Figures and Suppl Tables for "RNA splicing alterations induce a cellular stress response associated with poor prognosis in AML": Supplementary Material Anande et al BioRxiv.docx

**Patient cohorts**

Non-M3 AML patients from the TCGA and Clinseq cohorts were included if they received intensive induction chemotherapy, were devoid of splicing factor mutations, and had comprehensive clinical information (TCGA; n=104, Clinseq; n=184). For the TCGA-AML discovery cohort, RNA-seq data and associated mutational and clinical data were retrieved (in 2017) from the NCI data portal ([https://portal.gdc.cancer.gov](https://portal.gdc.cancer.gov/" \t "_blank)) and the study publication ^1^.  The Clinseq-AML cohort was treated in Sweden between 1997 and 2014 with intensive induction chemotherapy. The bone marrow or peripheral blood samples for these patients were obtained at the time of diagnosis and sequenced for their transcriptome and a panel of frequently mutated genes ^2^. Details about sample preparation, sequencing coverage and data access have previously been published ^2^. Clinical data were retrieved from the Swedish Acute Leukemia Registry (SALR) ^3^ or from patient records. Transcriptomic data from the Beat-AML cohort was used to validate significance of the splicing signature. Non-M3 AML patients who received intensive induction chemotherapy and had available survival and gene expression data with informative reads across signature events were included in the analysis (n=85). Data from the AMLCG group ^4^ was also considered for validation purposes but excluded due to the lack of informative reads across all signature events.

**RNA-seq data analyses**

Data pre-processing

Quality control checks were performed on raw RNA-seq data using FastQC (v 0.11.5). Adapter contamination and low-quality sequences in Clinseq-AML samples were removed by FastxToolkit. TCGA raw data was found to be clean, hence no filtering was required.  Quality filtered data were then processed for uniform read length, as splicing algorithms/tools need all reads to be of uniform length across all samples.

****For detailed methods, please contact Dr. Ashwin Unnikrishnan :** **

**Supplementary Figure Legends**

**Supplementary Figure 1.** (A) Distribution of differentially spliced events identified comparing ELN^Fav^ and ELN^Adv^ in the TCGA (left panel) and Clinseq (right panel) cohorts. SE, skipped exons. RI, retained introns. MXE, mutually exclusive exons. A5’SS, alternative 5’ splice sites. A3’SS, alternative 3’ splice sites. (B) Sashimi plot of a representative alternative 3’ splice site usage event in the *SSH3* gene in the TCGA data. Sequencing reads indicate usage of an alternate 3’ splice site in ELN^Adv^ patients (representative patients: #2820, #2838, red tracks) compared to ELN^Fav^ patients (representative patients: #2835, #2818, orange tracks). Lines connecting each exon represent splice junctions and numbers on each line represent number of supporting RNA-seq reads. (C) Sashimi plot of a representative alternative 5’ splice site usage event in the *ULK3* gene in the TCGA data. Representation as in (B). (D) ClueGO network interaction diagram of the 222 commonly spliced genes, with nodes representing functional pathways and edges indicating functional connections. Related to Figure 1.

**Supplementary Figure 2.** (A) LOGO analyses of splice acceptor sites of exons differentially retained (left) or skipped (right) in ELN^Adv^ patients. Boxed region indicates the +1 position. (B-D) Motif scanning analyses for PABPC1 (B), RBM46 (C) and SRSF3 (D) binding sites across a meta-exon generated from the differentially spliced events, with arrows indicating peaks of significant over-enrichment. Motif enrichment scores (left axes) and P values (right axes) are shown. The dashed lines indicate scores of skipped (red) and retained (blue) exons, while the black solid line indicates that of a background score from all non-differentially spliced exons. The green horizontal lines are set at p=0.05. (E) Interaction network indicating validated protein-protein interactions (edges) between the differentially spliced, with predicted functional impairment, splicing genes (nodes). Related to Figure 3.

**Supplementary Figure 3.** (A) Harrell’s C-index of risk classification by ELN, Splicing signature or LSC17 of TCGA-AML (left) or Clinseq-AML (right) patients. (B) TCGA-AML patients classified initially by ELN (left panel) and re-classified by the Splicing Signature (right panel). Sankey flow diagrams (middle panel) illustrate the redistribution of patients, with the widths of the lines proportional to numbers of patients redistributed (number also denoted). P-values were computed using Log-rank (Mantel-Cox) test. (C) TCGA-AML patients classified initially by LSC17 (left panel) and re-classified by the Splicing Signature (right panel). Representation similar to (A). (D) Clinseq-AML patients classified initially by ELN (left panel) and re-classified by the Splicing Signature (right panel). Representation similar to (A). (E) Clinseq-AML patients classified initially by LSC17 (left panel) and re-classified by the Splicing Signature (right panel). Representation similar to (A). (F-G) Harrell’s C-indices comparing risk classification by ELN, ELN + Splicing Signature, LSC17 or LSC17+ Splicing Signature of TCGA-AML (F) or Clinseq-AML (G) patients.

**Supplementary Tables**

**Supplementary Table 1**: Patient characteristics

****For a complete set of Supplementary Tables, please contact Dr. Ashwin Unnikrishnan:** ******
