## Supplementary figures and images for "RNA splicing alterations induce a cellular stress response associated with poor prognosis in AML"

### Supplementary Figure 1.pdf

Supplementary Figure 1

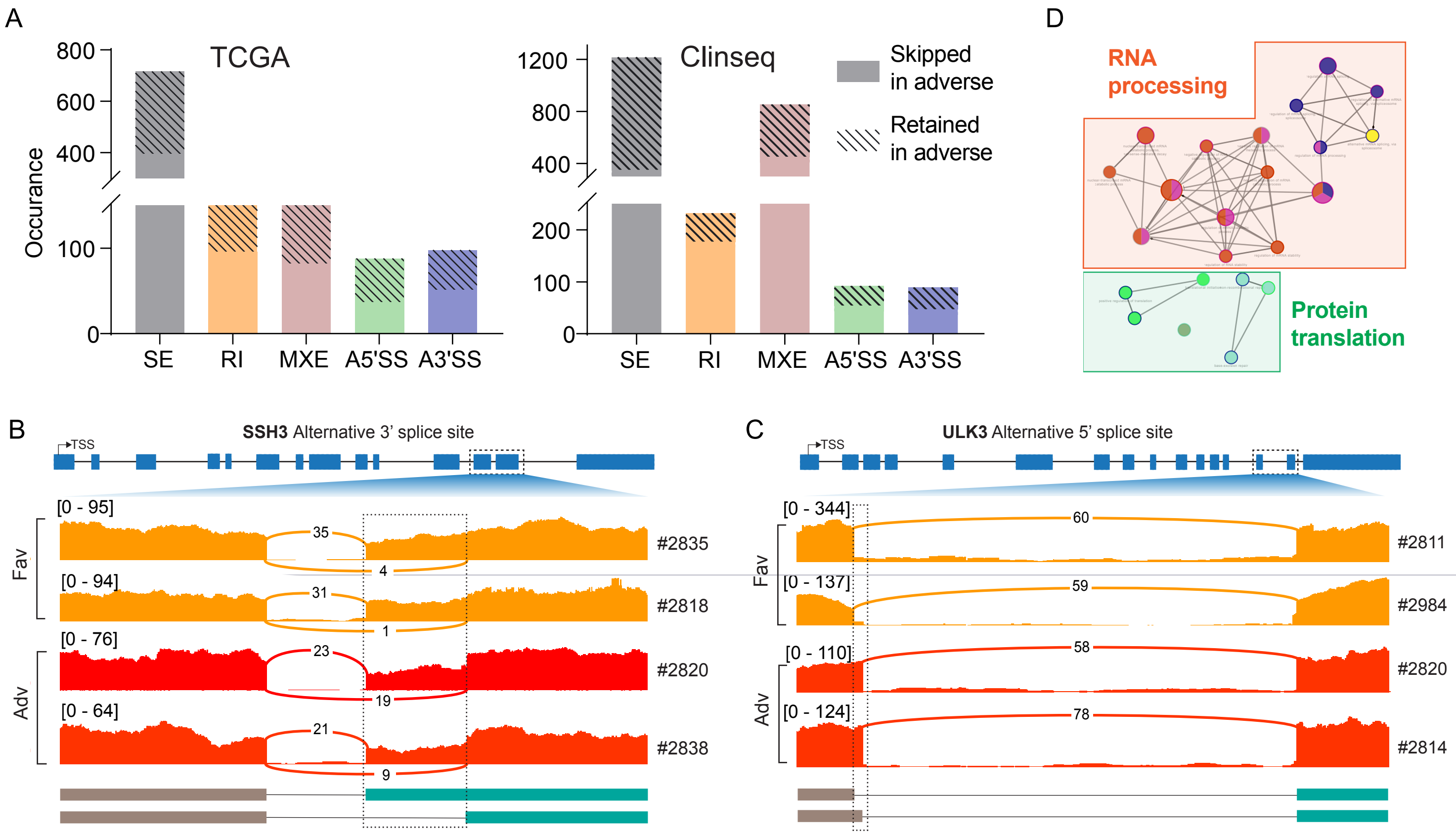

### Supplementary Figure 2.pdf

Supplemental Figure 2

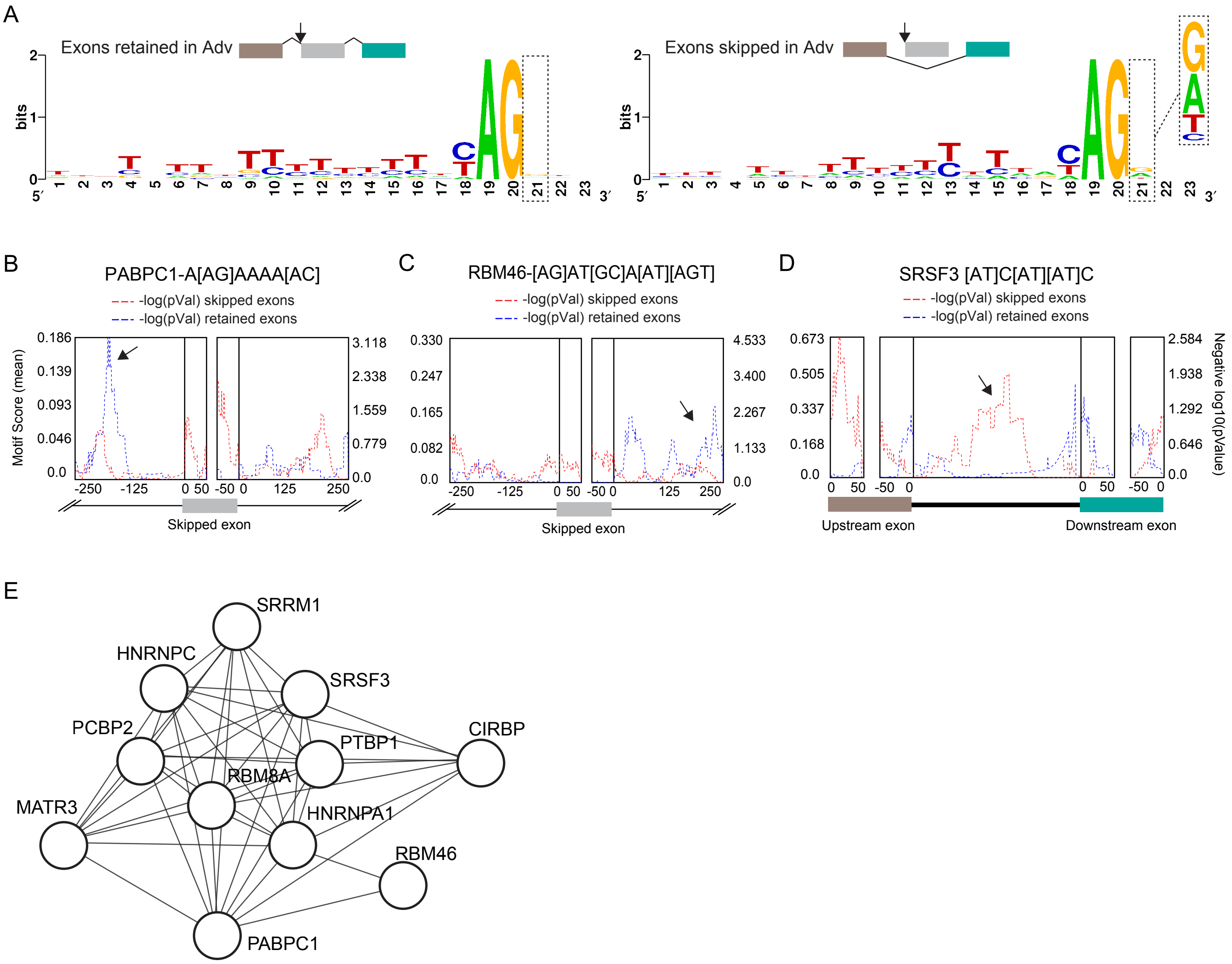

### Supplementary Figure 3.pdf

Supplemental figure 3

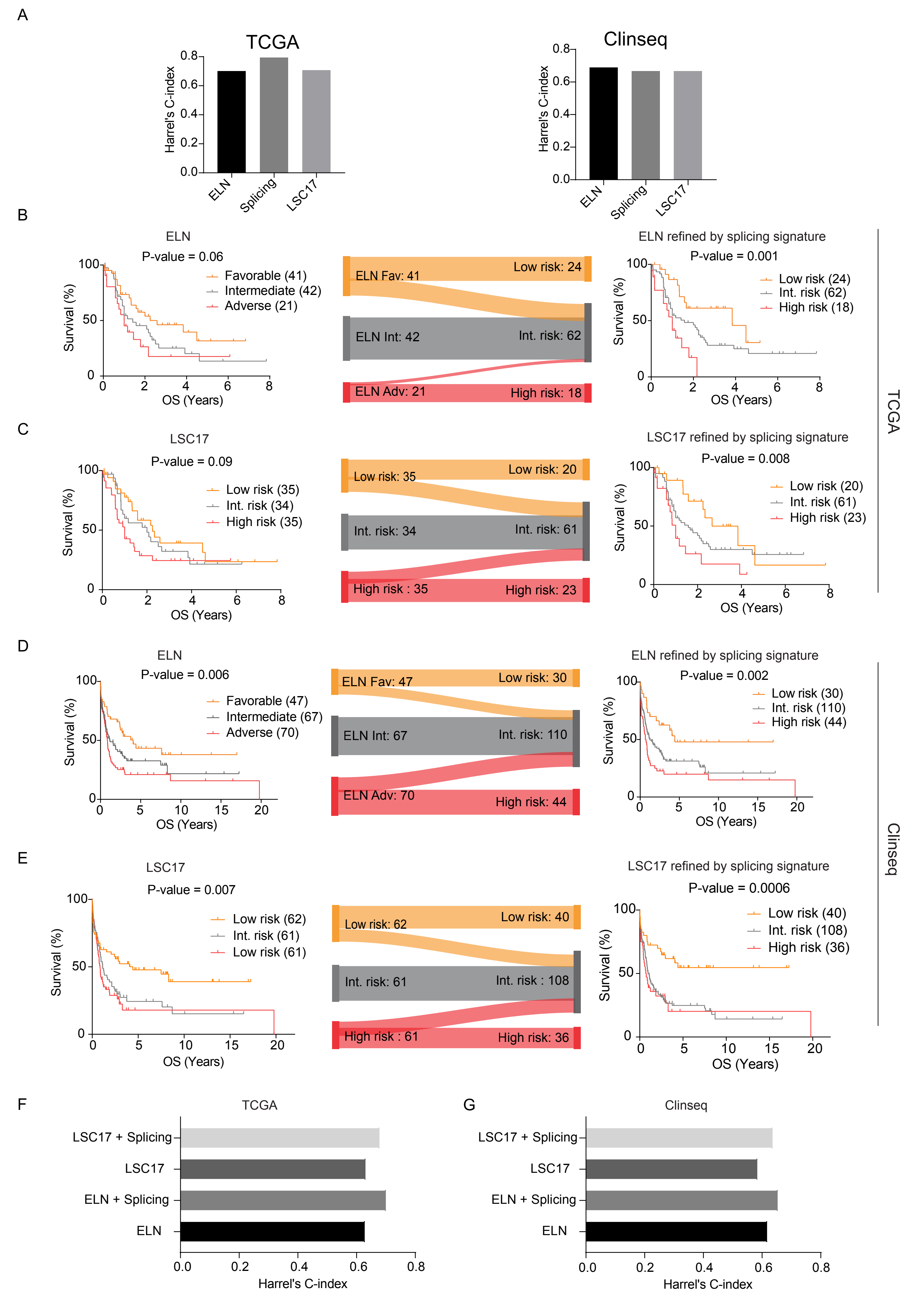
